# Apollo 3: Multi-Species Genome Curation

**DOI:** 10.64898/2026.06.08.730970

**Authors:** Garrett J Stevens, Kyösti Sutinen, Dario Beraldi, Shashank Budhanuru Ramaraju, Colin Diesh, William Haese-Hill, Peter Xie, Caroline Bridge, Adrien Morison, Angel Leung, Ulrike Böhme, Scott Cain, Nathan Dunn, Toby Hunt, Jane E Loveland, Adam Frankish, Alexie Papanicolau, Stefano Giorgetti, Jon Keatley, Bethany Flint, Lincoln Stein, Robert Buels, Matt Berriman, Ian Holmes

## Abstract

We present Apollo 3, a new manual genome annotation tool that integrates with the JBrowse 2 genome browser. Its functionality is inspired by existing manual genome annotation tools such as Apollo, Artemis, and Otter, combining features from those tools with an updated and more scalable architecture and technology stack. It allows simultaneous editing of multiple genomes, including visualization of synteny to inform those annotations. Apollo 3 can be used as a standalone annotation editor, or it can be installed on a server and used collaboratively.

## Introduction

Genome annotation is the process of taking the DNA sequence of an organism and creating analyses and interpretations to understand its biological significance (1). While automated methods exist for these analyses to determine both structure and function of the genomic sequence, they are not yet able to capture all information available in the genome (2). Manual curation and refinement by expert biocurators, especially when supported by cross-species comparison data, remains indispensable for high-quality, accurate annotations (3).

For more than two decades, Apollo (4) and Artemis (5) have been the two leading freely accessible tools to manually review and edit annotations. Artemis, a desktop application, has been used to annotate and analyze relatively small proand eukaryotic microbial genomes, often making use of intuitive synteny views available in its companion Artemis Comparison Tool (ACT) (6). Apollo, like Artemis, started as a desktop application (7), but was subsequently redesigned as a web-based application (8). Apollo used the JBrowse 1 genome browser (9) as a front end and offered improved browsing and editing functionality, coupled with simultaneous, collaborative annotation.

Other manual annotation tools have been influential in the biocuration community. A notable example is Otter (10), which has been primarily used by the team responsible for the Ensembl/GENCODE human and mouse reference gene annotation. They have used Otter to annotate more than 300,000 transcripts (11), and Otter’s unique interfaces include a rich transcript editor not found in other manual annotation tools.

While powerful, these tools have limitations. Artemis struggles with large genomes, is difficult to install, and lacks real-time collaborative updates. Apollo lacks the synteny-based visualizations found in Artemis and ACT. Otter remains a desktop-bound tool tied to specific infrastructure. As none are actively developed, these limitations will persist.

Here we present Apollo 3, a new manual genome annotation tool. Apollo 3 provides collaborative annotation capabilities like Apollo. Apollo 3 also supports annotation editing within synteny visualizations, similar to those in ACT, by integrating with JBrowse 2 (12). Apollo 3 was designed with extensive input from users of Apollo, Artemis, and Otter, so that it can fulfill the needs of annotators and overcome the limitations of the existing tools. Apollo 3 provides a new, flexible option for manual genome annotation.

### Design and Implementation

Apollo 3 includes three separate software products. First is the Apollo JBrowse Plugin, which is a JBrowse 2 plugin and provides the graphical user interface for Apollo 3. Next is the Apollo Collaboration Server, which provides the backend that supports collaborative annotation and interfaces with the database. Finally, the Apollo Command Line Interface (Apollo CLI) allows for programmatic interaction with the Apollo Collaboration Server.

The Apollo JBrowse Plugin can be installed without an Apollo Collaboration Server, allowing local single-user editing of genome annotations. Users will often also want to install the Apollo Collaboration Server in order to use its collaborative genome editing capabilities. Fig 1 shows the differences between these installations.

**Fig 1:**
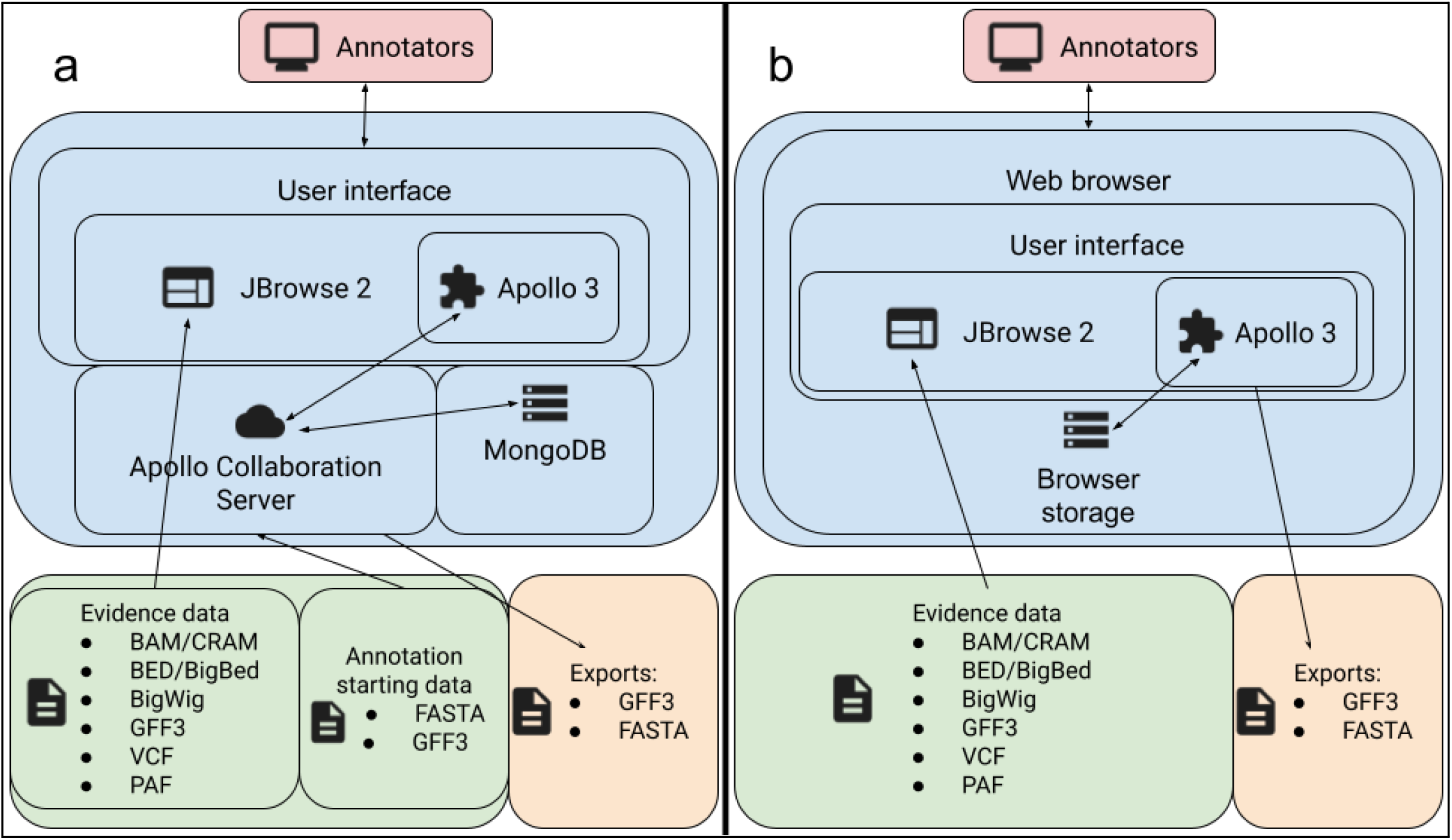
Apollo architecture diagram. (a) shows the architecture for Apollo 3 when used for collaborative editing, and (b) shows the architecture for Apollo 3 when used for local single-user editing. In both scenarios, Apollo components (in the central blue box) provide a way for annotators (red box) to edit genome annotations. For the collaborative case, annotation data is stored in a MongoDB database that is managed by the Apollo Collaboration Server. For the local case, annotation data is stored in the user’s browser storage. Input data (green box) includes evidence data that is viewed with JBrowse 2. Annotation data can be exported from either store in GFF3 and FASTA format (orange box).

### Apollo 3 has deep JBrowse 2 integration

JBrowse 2’s plugin architecture allows many ways for plugins to integrate with its functionality. Apollo 3 provides its entire user interface through these JBrowse 2 integrations, adding custom sidebars, tracks, and other displays. This is unlike the previous Apollo, which added a separate external user interface around JBrowse. The direct integration of Apollo 3 with JBrowse 2 allows those already familiar with JBrowse 2 to easily explore Apollo 3’s features.

As a JBrowse 2 plugin, Apollo 3 seamlessly utilizes its view types, including synteny views. These views can show comparisons between two or more species in a pairwise manner. These different comparisons can be quickly turned on and off in a similar way to linear evidence tracks. A user can, for example, use a DNA sequence alignment like that from minimap2 (13) to view broad alignment blocks. They can then switch to a translated nucleotide alignment like that from TBLASTX (14) for a more granular view of the relationships.

These synteny views can be used to compare related regions between genomes and to inform annotations of a genome by comparison with other related genomes whose annotations are already of high quality. For instance, conservation of synteny can extend beyond the coordinates of an existing gene model when compared to another species, allowing a gene model to be expanded.

### Apollo tracks provide a graphical interface for data

The main points of user interaction with Apollo 3 are two genome browser tracks within JBrowse 2: the annotations track, which Apollo 3 automatically adds for all of its assemblies, and the JBrowse 2 reference sequence track, which Apollo 3 customizes.

The primary area for annotation visualization in Apollo is the **annotations track**. Data can be displayed in this track in two ways, and users can switch between these displays in the track menu. In the **default display**, each feature takes up one or more rows, depending on its subfeatures. For example, gene features are drawn with a specific glyph that shows each transcript on a different row. This type of layout is common to many JBrowse 2 tracks. In the **six-frame display**, inspired by the annotation layout in Artemis, exons of a gene model are shown on different rows according to their reading frames (three forward or three reverse), with other non-exon features shown on two extra rows in the middle (for forward and reverse). In both displays, users can click and drag the boundaries of features to edit their coordinates. They can also right-click features to access more actions for that feature.

The Apollo track menu exposes several additional display options. In particular, under the “Appearance” submenu, users can choose to display a summary table of annotations, with subfeatures shown beneath their parent features. This table view allows users to view and edit details, such as coordinates and other attributes, about several features at the same time.

Apollo 3 also customizes the **JBrowse 2 reference sequence track**. It colors reading frames in its translation rows to match those shown in the annotations track and displays highlights that match selected features in the annotation track.

### Sidebars show details of on-screen data

Apollo 3 also adds various sidebars to JBrowse 2. These display details while not obscuring the tracks. The sidebars can be used to view and edit the details for a single Apollo feature annotation and are generally accessible by clicking on the desired feature in the track.

The design of these sidebars, especially in the case of gene or pseudogene transcripts (seen in Fig 2), is heavily influenced by Otter, including the feature coordinate display and a shortcut for trimming the coding region to match the displayed start and stop codons.

**Fig 2:**
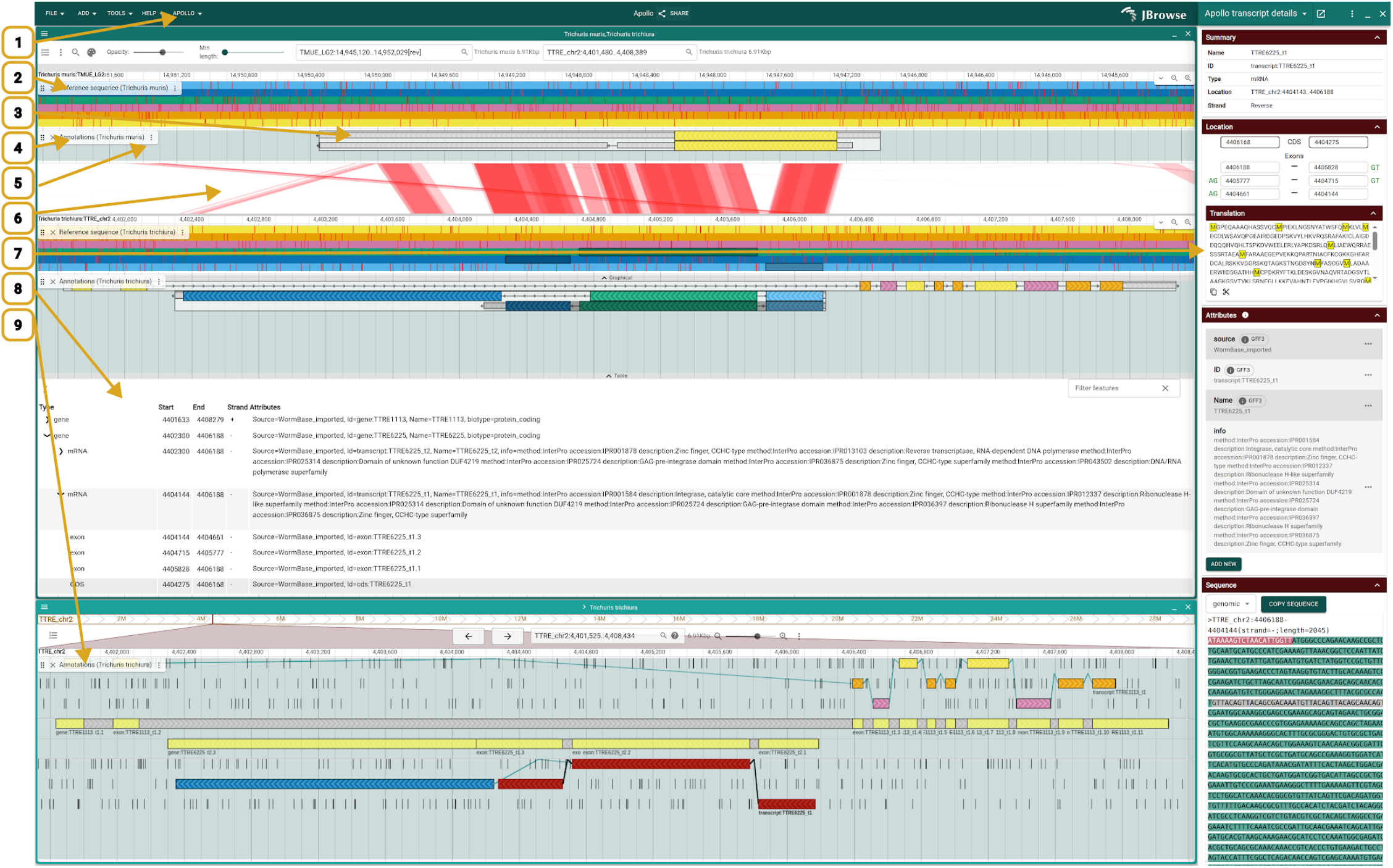
Key user interface elements of Apollo being used to annotate two related Trichuris species. (1) The top-level Apollo menu button. (2) A reference sequence track using the Apollo reference sequence display. (3) A glyph showing a gene with two transcripts in the default display. (4) An annotations track, shown with the default display. (5) A track menu button, used to access options regarding how the track is displayed. (6) A synteny track showing relationships between the species in the views above and below it. (7) An Apollo sidebar displaying information about a selected transcript, including the shortcut (the scissors icon) for trimming the coding region. (8) An Apollo annotation table display. (9) An annotations track in a separate view with the six-frame display. This view shows only the bottom species of the two in the top view.

### Apollo Collaboration Server enables shared, collaborative annotation editing

The Apollo Collaboration Server, used in collaborative installations, is a backend server process that facilitates storage, editing, retrieval, and real-time updates of annotations, as well as functions like user and login management.

The Apollo Collaboration Server uses MongoDB as its database. MongoDB is a fast, document-oriented, operational database that allows developers to store flexible JSON-like documents. The JSON format stored in MongoDB is also the same format that the front-end libraries use, removing the need to convert data representation between the back and front ends. MongoDB also provides text and genome range indexing of documents.

In particular this genomic range indexing needs to be performant in Apollo 3. Such queries form the basis for selecting the annotations to display on screen. The previous version of Apollo (referred to in this section as Apollo 2) loaded all annotations from a whole chromosome when viewing them in the application, instead of just the visible range; consequently, users who bulk loaded many annotations on a single chromosome experienced a noticeable drop in performance.

To test performance of both the initial indexing and subsequent query steps, we bulk-loaded all WormBase gene annotations from a single chromosome into Apollo 3 and Apollo 2. When we viewed a 50kbp region in the app, the time to retrieve the annotations in Apollo 3 was less than a quarter second, while the time for Apollo 2 was over 40 seconds.

This performance benefit comes at the cost of slightly increased time to initially load a genome onto the server in Apollo 3 vs Apollo 2. Where loading a genome with 1,000 chromosomes and contigs might take around a second in Apollo 2, it might take four seconds in Apollo 3. However, this is a one-time and probably tolerable cost: loading a given genome only needs to happen once (and is a task performed by an administrator), whereas querying will happen many times over the lifetime of a genome. We therefore find this tradeoff to be reasonable.

### Apollo Command Line Interface allows programmatic interactions with the Apollo Collaboration Server

Most users will interact with Apollo through the Apollo JBrowse Plugin’s user interface. For Apollo administrators, however, there is often a need to automate back-end tasks, such as adding a new assembly or importing new annotations. For these operations, we provide the Apollo Command Line Interface (CLI), which provides an interface for carrying out various Apollo operations via a terminal.

## Results

A well-designed application is more than a collection of features: these features must work together in a way that helps the user towards their goal. In this section, we present examples of Apollo 3 being used to curate genome annotations in different types of projects.

### Multi-genome curation

Here we explore the usage of Apollo 3 in combination with the comparative genomics capabilities of JBrowse 2 to improve the annotation set for the nematode *Trichuris trichiura*. To assist, we have annotations and synteny information for the related *Trichuris* species *T. suis* and *T. muris*.

Using a list of orthologs obtained for the three species, we found a pair of orthologous genes identified as WBGene00294573 in *T. muris* and D918_05905 in *T. suis* that did not have an ortholog identified in *T. trichiura*. Examining D918_05905 in JBrowse, we found a relationship identified by minimap2 between the surrounding area of the *T. suis* genome and a region of the *T. Trichiura* genome. We then ran TBLASTX between that region in *T. trichiura* and the region surrounding D918_05905 as well as the region surrounding WBGene00294573 and visualized those results in JBrowse 2 as well, confirming the relationships among the regions. Because the region was on the opposite strand in *T. trichiura* as compared to *T. suis* and *T. muris*, the ability of JBrowse 2 to reverse its view was critical to being able to visually confirm these relationships.

We then also visualized transcriptomic RNA-Seq reads in JBrowse that supported the existence of a gene in the identified region of the *T. Trichiura* genome. We used one of the reads to create a new gene and transcript in Apollo using the “Create Apollo annotation” menu item from the read’s context menu.

Upon examining the newly generated transcript, it became apparent that there was no reading frame that contained an appropriate translation start site for the transcript, but using the Apollo Reference Sequence Display, we identified that if the transcript were extended an extra two base pairs in the 5′ direction, it would then include such a site. After extending the coordinates of the gene and transcript in Apollo 3, we added a CDS region with the “Add coding sequence” option from the right-click menu.

We then obtained the resulting protein sequence for the new transcript from the transcript editor sidebar, as well as the sequences for the identified *T. suis* and *T. muris* genes. We compared these sequences and found that the new transcript created for *T. trichiura* codes for a protein that has 82% identity with the protein from D918_05905 in *T. suis* and 75% identity with the protein from WBGene00294573 in *T. muris*.

**Fig 3:**
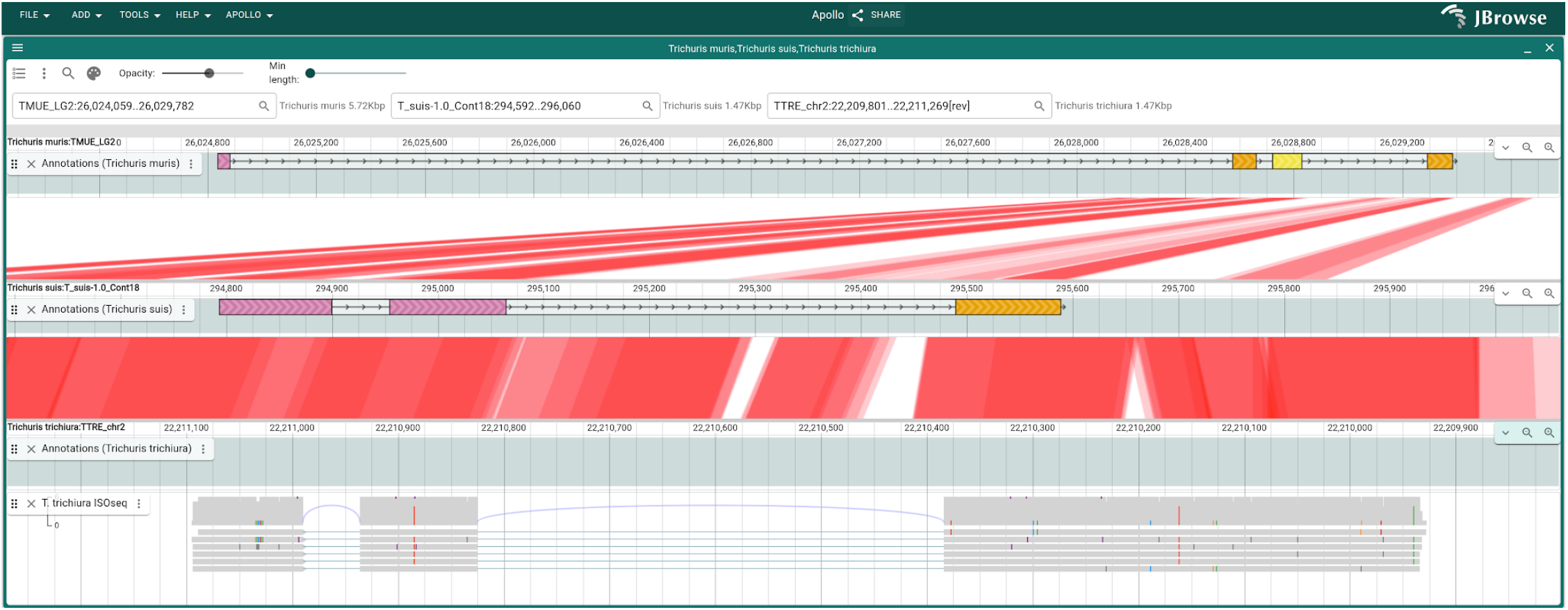
JBrowse 2 and Apollo 3 showing evidence for a missing gene in T. trichiura, with supporting evidence from two related Trichuris species. Three related Trichuris species Three different Trichuris species are shown in JBrowse 2’s linear synteny view. The Apollo annotations track is open for each species, and pairwise alignment obtained via TBLASTX is shown between the T. muris and T. suis pair as well as the T. suis and T. trichiura pair. The evidence indicates that T. trichiura is missing a gene annotation at the shown location. This is supported by an RNA-seq track that is also shown for T. trichiura. The gene created based on this evidence codes for a protein that has 82% identity with a protein from the gene shown in T. suis and 75% identity with the protein from the gene shown in T. muris.

### Single-genome collaborative curation

In this example, curators from the Ensembl-HAVANA team at EMBL-EBI collaboratively edit reference annotations for human genes as part of the GENCODE project (11). Apollo 3’s collaborative editing makes updates immediately visible, helping users avoid conflicting edits when working on the same or nearby annotations.

During the annotation process these curators use many evidence tracks to evaluate and update annotations. Some tracks are used to visually guide annotations, and some tracks have additional Apollo 3 integrations available. These include JBrowse 2 feature tracks, typically used with GFF3 or BED files that have gene or transcript models. Curators copy genes and transcripts from these tracks into the Apollo 3 annotations. A copied transcript can be integrated into an existing gene or have a new gene automatically generated for it. Similarly, curators can copy reads from JBrowse 2 alignments tracks, especially long reads from transcriptomic data, into Apollo 3 new genes or transcripts as well.

The HAVANA team has also developed their own plugins to customize Apollo 3. For example, to augment native GFF3 export ability, they have added the ability to export annotations to their own existing database.

**Fig 4:**
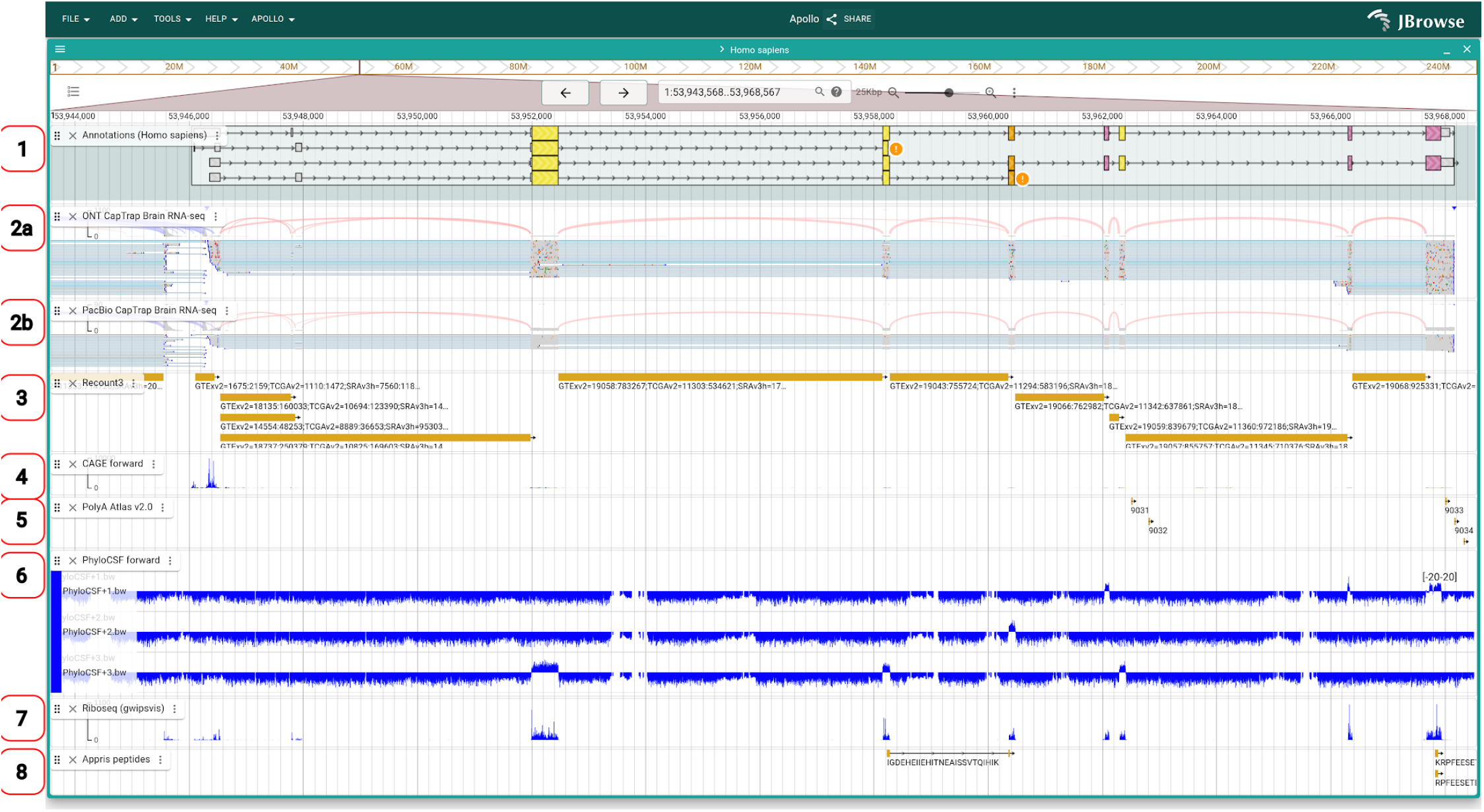
Apollo 3 with multiple evidence tracks. An example of a gene and evidence tracks that support its model. See Table 1 for a description of the evidence tracks. The tracks are (1) Apollo annotation track (2) LRGASP RNA-seq (3) Recount3 RNA-seq (4) CAGE TSSs (5) PolyASite TSEs (6) PhyloCSF conserved regions (7) GWIPS-viz Ribo-seq (8) APPRIS

**Table 1.** Example evidence tracks and how they are used for curating annotations.

| Track | Type | Description |
| --- | --- | --- |
| LRGASP<br>RNA-seq | BAM/<br>CRAM | In addition to the models provided by LRGASP(15), the raw reads from both Oxford Nanopore Technologies and PacBio sequencing are used to evaluate structural annotations. |
| Recount3<br>RNA-seq | BED | Intron position data based on short-read data from Recount3 (16) is used to provide orthogonal support for transcript structure. |
| CAGE TSSs | BigWig | Cap analysis of gene expression (CAGE) data from the FANTOM5 (17) project is used to locate transcription start sites (TSSs). |
| PolyASite TSEs | BED | PolyASite V3.0 (18) single-cell RNA-sequencing data is used to infer Transcription End Sites (TSEs) from polyadenylation sites. |
| PhyloCSF<br>conserved regions | BigWig | PhyloCSF(19), a method that detects evolutionary signatures of protein-coding regions using multi-species genome alignments, is used to evaluate the coding potential, or lack thereof, of a transcript. |
| GWIPS-viz<br>Ribo-seq | BigWig | The aggregation of multiple Ribo-seq experiments as available via the GWIPS-viz browser (20) is used to profile ribosomal data and evaluate the coding potential, or lack thereof, of a transcript. |
| APPRIS | BigBed | APPRIS project (20,21) mass spectrometry peptide evidence supports annotated coding transcripts to evaluate the coding potential, or lack thereof, of a transcript. |

### Single-genome single-user curation

In this example, we curate annotations from the genome of *Onchocerca volvulus* (∼100 Mb) in JBrowse 2 Desktop. We use the local editing capabilities of Apollo 3, without using the Apollo Collaboration Server. After opening the genome using FASTA files from a local hard drive and installing the Apollo JBrowse Plugin from the JBrowse 2 plugin store, an Apollo track was automatically added for the organism. We then loaded features into the track from a GFF3 file and enabled the six-frame view. This six-frame view, based on the default annotation view used in Artemis, provides intuitive visualization of the consequences of exon boundary edits (i.e. dynamically shifting downstream exons into different reading frames), which was useful for improving the existing annotations.

**Fig 5:**
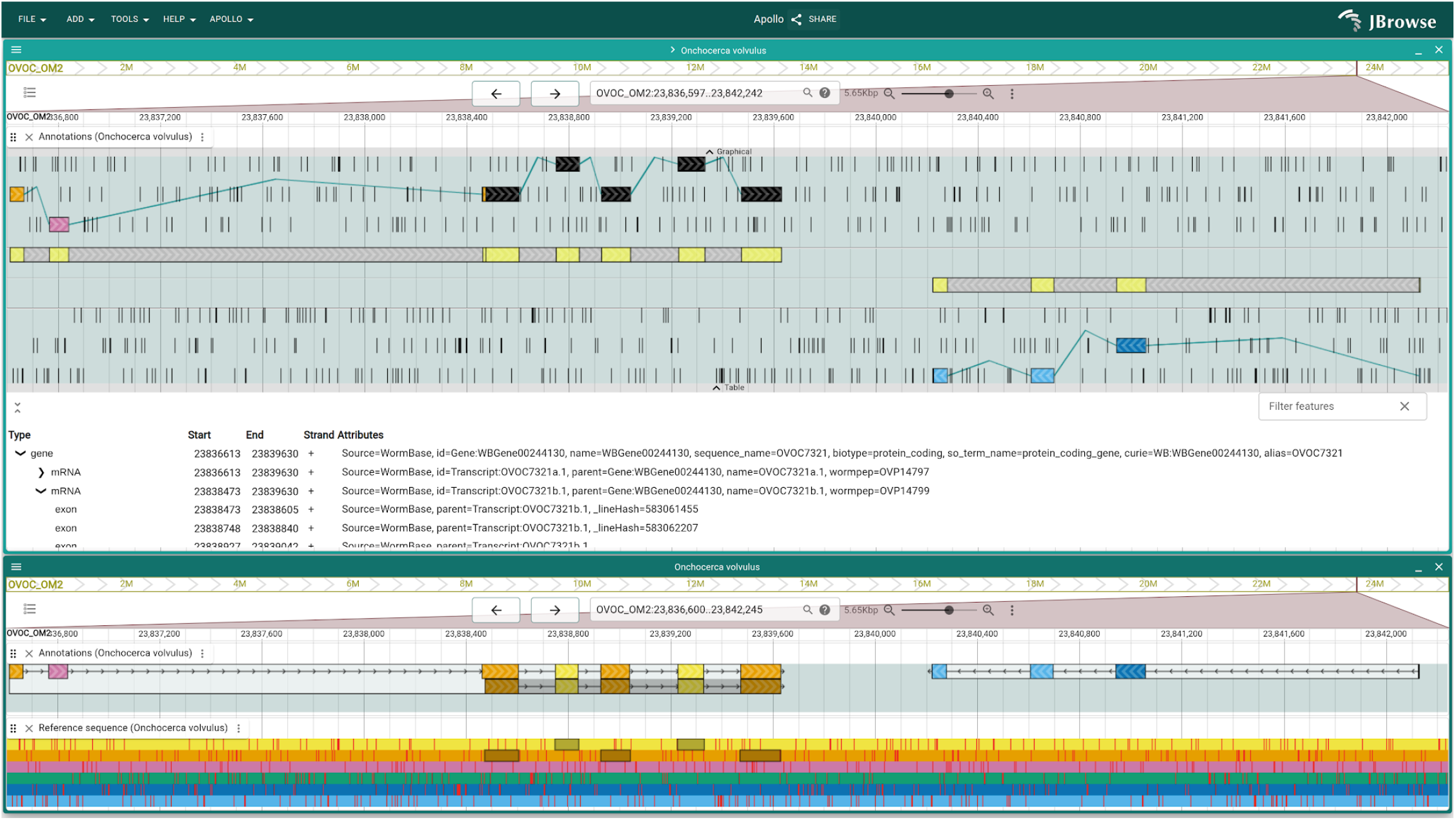
Onchocerca volvulus being annotated with Apollo 3 on JBrowse 2 Desktop. Shown are two Apollo annotation tracks with standard and six-frame display, showing gene OVOC7321 from the O. volvulus reference genome, with two transcripts. The tracks are shown in two separate views. Transcript OVOC7321b.1 is highlighted clearly in both the six-frame display and default display when hovering over the feature row in the table view. Also note the adjacent gene OVOC7322, which illustrates the distinction between genes on oppsosite strands.

## Availability and Future Directions

### Availability

The source code of Apollo 3 is available on GitHub (22) and is published under the Apache 2.0 license. An archival copy of the source code is available on Zenodo(23). Installation guides and documentation are available at the project home page (24). A list of live demos associated with the figures in this paper is also available on the project’s website (25).

The Apollo Collaboration Server and the Apollo CLI are available as Docker images on the GitHub Container Registry: ghcr.io/gmod/apollo-collaboration-server and ghcr.io/gmod/apollo-cli. The Apollo CLI can also be installed from the NPM registry under the name @apollo-annotation/cli.

### Future Directions

#### AI anticipation of annotator actions

The extensive plugin capabilities of JBrowse 2 allow many different types of plugin integrations, including plugins that interact with Apollo 3. One such possible plugin functionality is instrumenting JBrowse 2 to be able to track what users are looking at and what actions they take. These observations could then be used to train an AI model that could predict annotator actions, allowing Apollo 3 to anticipate and streamline workflows for annotators.

#### Liftover of annotations using synteny evidence

Synteny evidence can show relationships between genomes. These relationships could be used to lift over a gene model from one genome into another, estimating its most likely information based on one or more synteny datasets. Because this would likely not offer base-pair accuracy of the new gene’s position, other information such as the boundaries of features in supporting evidence tracks and calculation of open reading frames could also inform the liftover.

#### Integration with de novo gene finder

An alternative to lifting an existing gene over into a genome where that gene appears to be missing is to run a de novo gene finder to predict the gene’s structure. This gene finder could be run on the Apollo Collaboration Server for collaborative installations. To support standalone installations, we can port a gene finding algorithm to WebAssembly and run it for small regions directly in the browser.

## Acknowledgements

In addition to the JBrowse 2 developers who contributed directly to Apollo 3 and are authors on this paper, we would also like to thank Teresa De Jesus Martinez and Elliot Hershberg for their work on JBrowse 2 and its plugin ecosystem. We would also like to thank the following members of the Apollo community call group who provided expert feedback on Apollo planning: Chris Childers, Paul Wilkinson, Stavros Diamantakis, and David Starns.

